# A cell wall invertase controls nectar volume and sugar composition

**DOI:** 10.1101/2021.03.05.434125

**Authors:** Anzu Minami, Xiaojun Kang, Clay J. Carter

## Abstract

Flowering plants produce nectar to attract pollinators. The main nectar sugars are sucrose, glucose, and fructose, which can vary widely in ratio and concentration across species. *Brassica* spp. produce a hexose-dominant nectar (high in the monosaccharides glucose and fructose) with very low levels of the disaccharide sucrose. Cell wall invertases (CWINVs) catalyze the irreversible hydrolysis of sucrose into glucose and fructose in the apoplast. We found that *BrCWINV4A* is highly expressed in the nectaries of *Brassica rapa*. Moreover, a *brcwinv4a* null mutant has (1) greatly reduced cell wall invertase activity in the nectaries, and (2) produces a sucrose-rich nectar with little hexose content, but (3) with significantly less volume. These results were recapitulated via exogenous application of an invertase inhibitor to wild-type flowers. Honeybees prefer nectars with some sucrose, but wild-type *B. rapa* flowers were much more heavily visited than those of *brcwinv4a*, suggesting that the potentially attractive sucrose-rich nectar of *brcwinv4a* could not compensate for its low volume. These results cumulatively indicate that BrCWINV4A is not only essential for producing a hexose-rich nectar, but also support a model of nectar secretion in which its hydrolase activity is required for maintaining a high intracellular-to-extracellular sucrose ratio that facilitates the continuous export of sucrose into the apoplast via SWEET9. Extracellular hydrolysis of each sucrose into two hexoses by *BrCWINV4A* also likely creates the osmotic potential required for nectar droplet formation. In summary, modulation of CWINV activity can at least partially account for naturally occurring differences in nectar volume and sugar composition.

## INTRODUCTION

Flowering plants produce floral nectar to attract pollinators (Pacini and Nepi, 2007). These animal mutualists contribute to the sexual reproduction of ∼87% of all angiosperms (Ollerton et al., 2011), including 76% of all leading global crops that account for ca. 30% of all food production (Klein et al., 2007). Honeybees are the most economically and effective pollinators of most crops (Klein et al., 2007), thus, farmers often manage honeybee hives to ensure high pollination efficiency. For instance, honeybees dominate the flower-visiting insects of mustard (*Brassica rapa*), and this visitation increases seed yield (Atmowidi et al., 2007), even though mustard is partially self-compatible (Downey and Rimmer, 1993). The number of bee visits is also positively correlated with nectar volume, which is the main source of carbohydrates for honeybees (Silva and Dean, 2000). Plant nectars primarily contain varying ratios of the disaccharide sucrose and its hydrolysis products, the hexoses of glucose and fructose (Nicolson and Thornburg, 2007). Honeybees tend to prefer nectars containing moderate levels of sucrose to ones with glucose and fructose alone (Wykes, 1952). However, *Brassica* spp., produce hexose-dominant nectars with a very low amount of sucrose (Baker and Baker, 1983; Davis et al., 1998).

*Brassica* spp. (Brassicaceae) are major agricultural and horticultural crops, including vegetables (cabbage, turnip, broccoli, cauliflower, turnip, and Chinese cabbage, etc.) and oilseeds (canola and mustard) (Paterson et al., 2001; Johnston et al., 2005; Town et al., 2006). Besides Arabidopsis, members of the Brassicaceae are popular models for polyploid breeding and self-incompatibility studies (Paterson et al., 2001; Johnston et al., 2005; Town et al., 2006). Many kinds of Brassicaceae are cultivated all over the world, which are visited by many insects during the flowering season (Mishra and Sharma, 1988; Davis et al., 1996; Pierre et al., 1999; Masierowska, 2003; Bertazzini and Forlani, 2016).

Nectaries are the glands that produce nectar. *Brassica* flowers contain two non-equivalent sets of nectaries, lateral and median (Davis et al., 1986). Lateral nectaries produce the vast majority of nectar, whereas median nectaries produce little or no nectar (Davis et al., 1986; Davis et al., 1996; Davis et al., 1998). In the most accepted model of nectar production in the Brassicaceae, lateral nectaries actively synthesize sucrose from degraded starch at anthesis, which is unloaded to the apoplast via the plasma membrane-localized sucrose transporter SWEET9 (Lin et al., 2014). CELL WALL INVERTASE4 (CWINV4) subsequently converts the apoplastic sucrose into the hexoses glucose and fructose, thereby creating negative water potential and causing water flow out of the parenchymal cells (Ruhlmann et al., 2010). Finally, these soluble sugars, along with other nectar components, exit the nectary via open stomata (Roy et al., 2017).

Invertases (EC 3.2.1.26; β-fructofuranosidase) catalyze the irreversible hydrolysis of sucrose into glucose and fructose (Fig. 1A) and function in plant growth, development, and stress responses (Roitsch et al., 1995; Tymowska-Lalanne and Kreis, 1998; Roitsch and Ehness, 2000; Berger et al., 2007). Plants have three type of invertases: cell wall invertase (CWINV), vacuolar invertase (VINV) and cytosolic invertase (CytINV), which are divided into two subfamilies, acid invertases (CWINV and VINV) and neutral/alkaline invertases (CytINV) based on the optimum pH for activity (Tymowska-Lalanne and Kreis, 1998, 1998; Sturm, 1999; Winter and Huber, 2000). The *Arabidopsis thaliana* genome encodes six putative encoding cell wall invertases (CWINVs), which are all expressed in flowers, as well as various other tissues (Sherson et al., 2003).

**Figure 1.**
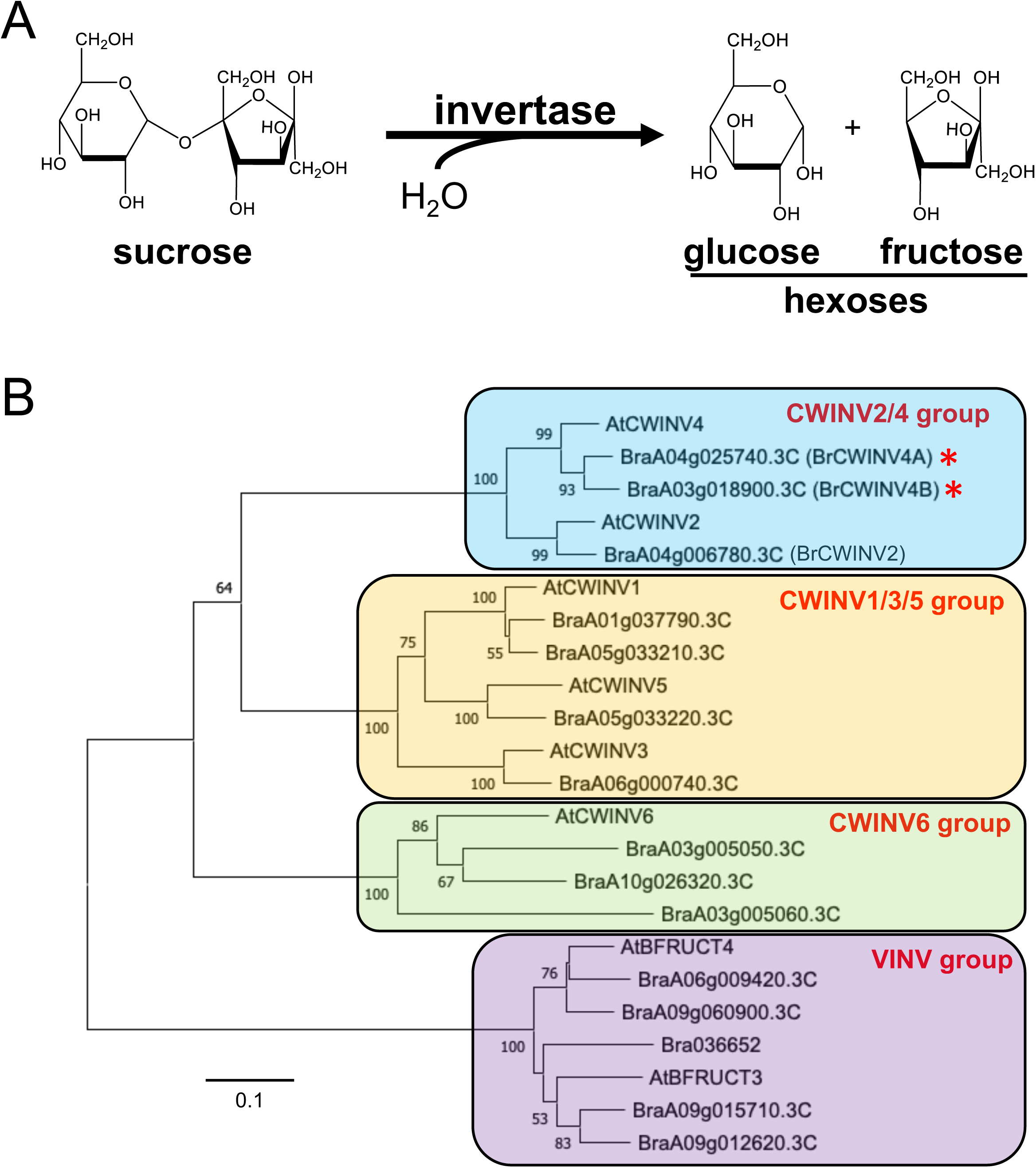
Phylogenic analysis of cell wall invertases in *Brassica rapa* and *Arabidopsis thaliana*. (A) Invertases catalyze the irreversible hydrolysis of sucrose to glucose and fructose. (B) Phylogenetic analysis of cell wall invertases in *Brassica rapa* and *Arabidopsis thaliana*. The scale bar represents 0.1 substitutions per sequence site. The numbers at nodes are bootstrap values from 100 replicates. CWINV, cell wall invertase; VINV, vacuolar invertase.

A role for invertases in controlling nectar sugar composition (i.e., sucrose- vs. hexose-rich) has long been postulated (Baker and Baker, 1983), but this hypothesis has been supported by limited experimental evidence. Since Arabidopsis *atcwinv4* mutants do not produce any nectar (Ruhlmann et al., 2010), it could not be definitively determined if AtCWINV4 activity regulates nectar sugar composition or maintains nectary sink status. Unlike Arabidopsis, *B. rapa* displays partial self-incompatibility and is dependent on pollinators to achieve full fecundity (Mishra and Sharma, 1988; Bommarco et al., 2012). Consequently, *B. rapa* flowers are much larger and more conspicuous than those of Arabidopsis. In this study, we investigated a potential role for cell wall invertases in the control of nectar composition and volume in *B. rapa*.

## RESULTS

Since *AtCWINV4* is required for nectar production in Arabidopsis (Ruhlmann et al., 2010), we looked for potential orthologs in *B. rapa*. Plant CWINVs belong to the glycoside hydrolase 32 family (GH32) together with VINVs, thus we obtained sequences of *B. rapa* GH32 genes from the Brassica Database (BRAD) and compared their predicted amino acid sequences with those of Arabidopsis CWINVs. This phylogenetic analysis shows that CWINVs both of *B. rapa* and Arabidopsis group into three distinct subfamilies, which are completely separate from a group of VINVs, including *betaFruct3* (*BFUCT3*) and *BFUCT4* (Fig. 1B). *AtCWINV4* and *AtCWINV2* fall into the same clade, with the *B. rapa* genome encoding two and one putative CWINVs that are similar to *AtCWINV4* and *AtCWINV2*, respectively. We named the two mostly closely related genes to *AtCWINV4* as *BrCWINV4A (BraA04g025740.3C*) and *BrCWINV4B (BraA03g018900.3C*). At the amino acid level, AtCWINV4 shares 86% identity with both *BrCWINV4A* and *BrCWINV4B*, whereas *BrCWINV4A* and *BrCWINV4B* have 89% identity with one another (Supplemental Fig. S1).

Since *AtCWINV4* displayed nectary-enriched expression (Ruhlmann et al., 2010), we examined the expression patterns of *BrCWINV4A*, *BrCWINV4B* and *BrCWINV2* by semi-quantitative RT-PCR (Fig. 2A, Supplemental Fig. S2A) and sequencing of PCR products (Supplemental Fig. S2B). Due to the high sequence similarity between *BrCWINV4A* and *BrCWINV4B*, it was difficult to design gene-specific primers for RT-PCR analysis, as both genes appeared to display nectary-enriched expression (Fig. 2A, Supplemental Fig. S2); however, sequencing of the RT-PCR products from nectary tissues indicated that *BrCWINV4A* is much more highly expressed in both the lateral (LN) and median nectaries (MN) than *BrCWINV4B* even when primers designed to be more specific for *BrCWINV4B* were used for RT-PCR (Supplemental Fig. S2B). Moreover, a prior EST analysis of *B. rapa* nectaries (Hampton et al., 2010) identified a putative orthologous cDNA (GenBank: GQ146458) to *AtCWINV4* that was highly represented in the EST library and much more similar to *BrCWINV4A* (99.5% identity) than *BrCWINV4B* (89% identity; Supplemental Fig. S3). Conversely, no nectary ESTs with similarly high identify to BrCWINV4B were identified in the prior study. Finally, *BrCWINV2* is highly expressed in stamens, but is only slightly expressed in nectaries (Fig. 2A).

**Figure 2.**
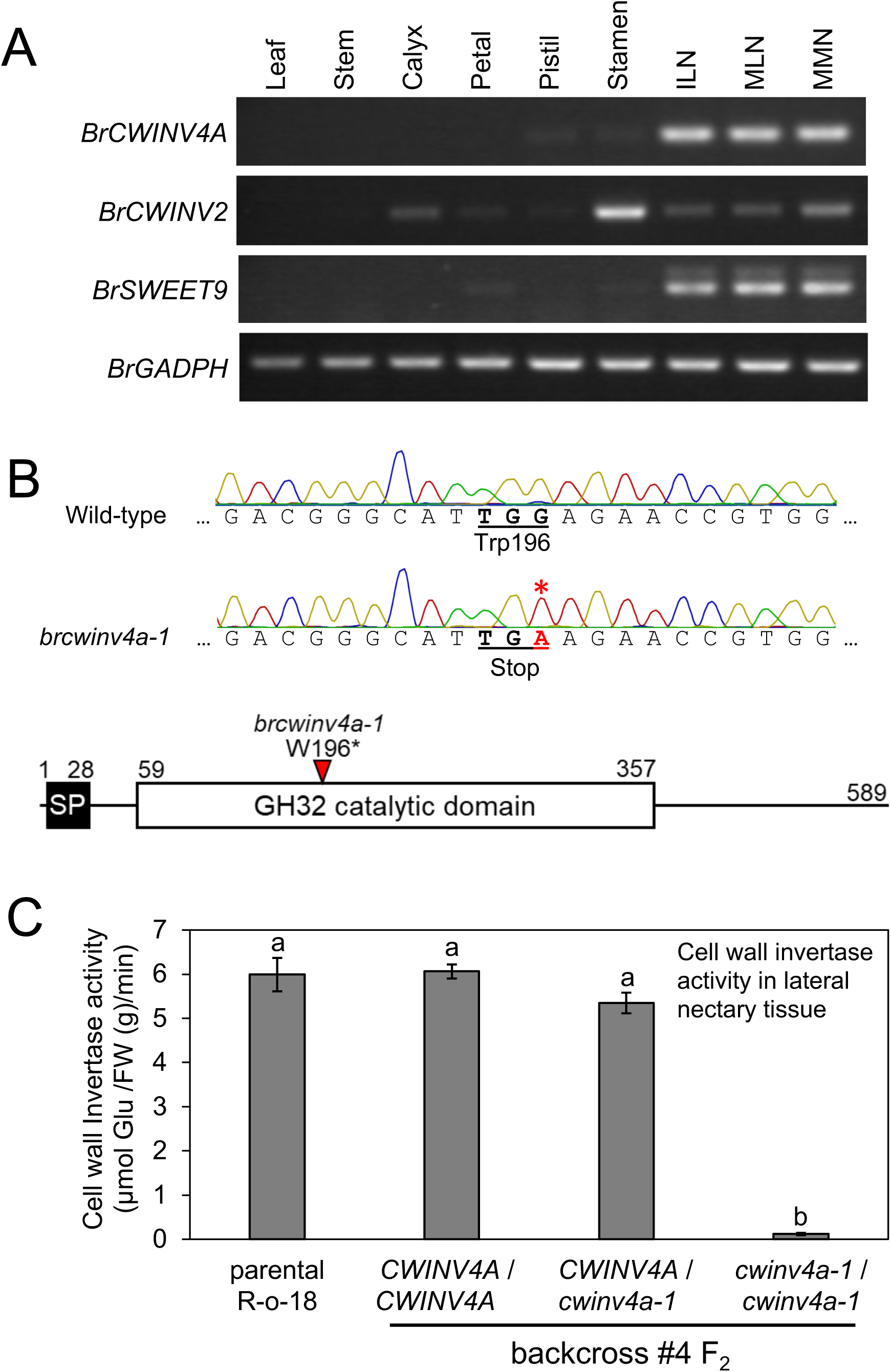
*BrCWINVA* is the main cell wall invertase in *Brassica rapa* nectaries. (A) RT-PCR analysis was performed using total RNA isolated from leaf, stamen, calyx, petal, pistil, stamen, immature lateral nectary [ILN, pre-anthesis, equivalent to Stage 12 in Arabidopsis (Smyth et al., 1990)] and mature lateral nectary [MLN, Stage 14–15 flower], and mature median nectary (MMN) of *B. rapa* wild-type (R-o-18). *BrGADPH* was used as a reference control and BrSWEET9 was used as a nectary-enriched expression control. (B) DNA sequencing chromatograms of wild-type and the TILLING mutant line Ji40447, which has a G-to-A point mutation at nucleotide at nucleotide +588 from the predicted start codon of the *BrCWINV4A* cDNA (indicated by red asterisk). This mutation creates a premature stop codon (TGA) in place of W196. The lower schematic indicates the location and nature of the mutation, as well as the predicted signal peptide (SP) for secretion from the cell. (C) Cell wall invertase activity in lateral nectaries after four backcrosses of Ji40447 with wild-type (R-o-18) followed by one-time self-pollination. Plants genotyping as wild-type, heterozygous, and homozygous for the W196* mutation are labeled as *CWINV4A*/*CWINV4A*, *CWINV4A*/*cwinv4a-1*, and *cwinv4a-1*/*cwinv4a-1* respectively. Cell wall fractions extracted from twenty lateral nectaries of Stage 14–15 flowers of wild type (R-o-18) and the BC4F_2_ lines were tested for the invertase activity. Bars represent averages of at least three independent biological and three technical replicates. Error bars show SE. Different letters indicate significant differences (P<0.05) according to the Tukey-Kramer test.

Since *BrCWINV4A* was determined to be the most likely direct ortholog of *AtCWINV4*, we obtained a TILLING (Targeting Induced Local Lesions in Genomes) mutant (Ji40447-b) for *BrCWINV4A* from the RevGenUK TILLING platform at the John Innes Centre (Stephenson et al., 2010). *BrCWINV4A* is predicted to encode a protein of 589 amino acids with a 28 amino acid signal peptide for secretion from the cell (Fig. 2B, Supplemental Fig. S1). Line Ji40447-b contains a G-to-A point mutation located at nucleotide +588 from the predicted start codon of the cDNA, converting Trp196 (TG<u>G</u>) into a stop codon (TG<u>A</u>; results in W196*). The premature stop codon exists in the N-terminal half of the predicted Glycosyl Hydrolase 32 (GH32) catalytic domain (Fig. 2B). This mutant line was subsequently backcrossed to the wild-type parent R-o-18 four times (BC4) prior to extensive phenotypic characterization (i.e., unless otherwise noted, phenotyping occurred in the BC4F_2_ or BC4F_3_ generation). The mutation was verified in each generation following backcrossing and self-pollination by sequencing of PCR products amplified from genomic DNA (and verified for the accompanying phenotype in homozygous mutants as described below). This line was renamed as *brcwinv4a-1* (Fig. 2B).

Brassicaceae flowers, including those of *B. rapa* and Arabidopsis, often have two pairs of lateral and median nectaries (Davis et al., 1986; Davis et al., 1995; Kram and Carter, 2009; Hampton et al., 2010). A vast majority of nectar is secreted from the lateral nectaries (LN) located interior of the base of a short stamens, whereas median nectaries (MN) form outside the base of long stamens and produce little or no nectar (Davis et al., 1986; Davis et al., 1995; Davis et al., 1996; Davis et al., 1998). Therefore, for functional analysis of *BrCWINV4A*, we analyzed cell wall invertase activity in the LN of the BC4F_2_ generation, which contained a mixture of homozygous (*brcwinv4a-1/brwinv4a-1*), heterozygous (*BrCWINV4A-1/brcwinv4a-1*) and wild-type (*BrCWINV4A-1/BrCWINV4A-1*) plants [i.e., derived from the original M_2_ TILLING mutant backcrossed with R-o-18 four times and self-pollinated in the F_1_ generation after each backcross (Fig. 2C)]. The cell wall invertase activity of the LN in wild-type (R-o-18) was 6.00 ± 0.38 µmol glucose g^-1^ FW min^-1^ (Fig. 2C). Among the BC4F_2_ generation, all plants with wild-type (*BrCWINV4A-1/BrCWINV4A-1*) and heterozygous (*BrCWINV4A-1/brcwinv4a-1*) alleles showed cell wall invertase activities similar to that of the wild-type parent (6.06 ± 0.16 and 5.34 ± 0.23 µmol glucose g^-1^ FW min^-1^, respectively). In contrast, the homozygous BC4F_2_ allele plants (*brcwinv4a-1/brwinv4a-1*) had almost no cell wall invertase activity in the lateral nectaries (0.12 ± 0.03 µmol glucose g-1 FW min-1). These results indicate that BrCWINV4A functions as the primary cell wall invertase in lateral nectaries and that *brcwinv4-1a* is likely a null mutant.

We next observed the nectar secretion patterns and droplet sizes from the lateral nectaries of homozygous *brcwinv4a-1* (BC4F_3_) plants (Fig. 3). Like Arabidopsis *atcwinv4* flowers (Ruhlmann et al., 2010), there was no observable difference in the structure of LNs and MNs between R-o-18 (parental) and *brcwinv4a-1* (Fig. 3A, Supplemental Fig. S4). All R-o-18 (wild-type) *B. rapa* flowers (100%) produced large nectar droplets from LNs after anthesis, with the two droplets on a single LN also sometimes merging to become one large droplet (Fig. 3A). Unlike Arabidopsis *atcwinv4*, ∼71% of *brcwinv4a-1* flowers produced nectar. Out of 202 *brcwinv4a-1* flowers from six individual plants, 59 (29.2%) did not produce observable nectar drops (Fig. 3A, right), 61 (30.2%) secreted only one small nectar drop, and 82 (40.6%) secreted two nectar droplets on an individual LN. While nectar droplet formation was not strictly quantified in the prior generations of segregating, backcrossed plants, all flowers from plants that genotyped as wild-type (*BrCWINV4A/BrCWINV4A*; N>20) and heterozygous (*BrCWINV4A/brcwinv4a-1;* N>15) produced large nectar droplets, whereas all homozygous plants (*brcwinv4a-1/brwinv4a-1*; N>20) displayed the same nectar secretion phenotype observed in the BC4F_2_ generation (i.e., either having no nectar or one or two small nectar droplets; Supplemental Fig. S4).

**Figure 3.**
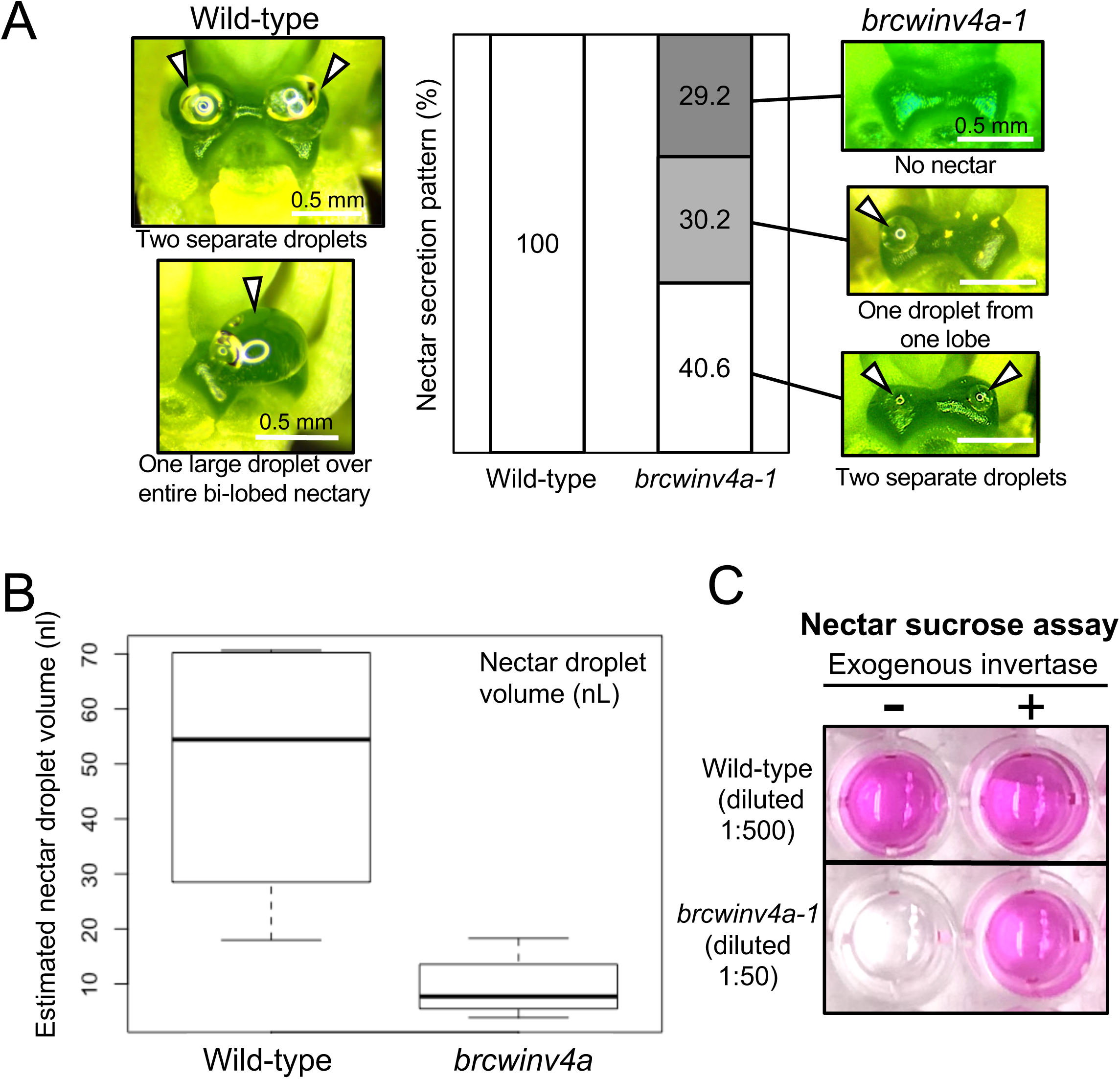
Nectar secretion patterns and droplet size in the lateral nectaries of wild-type and *brcwinv4a-1* homozygous plants. (A) Examples of nectar droplets from wild-type (R-o-18) (left) and *brcwinv4a-1* (right) flowers. Scale bars are 0.5 mm. All (100%) wild-type nectaries (left) produced large nectar droplets and the secretion pattern was observed as two individual droplets or a combined single large droplet. *brcwinv4a-1* nectaries were classified as producing either no nectar, one small droplet, or two small nectar droplets. The bar graph represents the percentage of different nectar secretion patterns from lateral nectaries (*N* = 202 flowers for both wild-type and *brcwinv4a-1* from six different plants). (B) Boxplot analysis of estimated nectar volume produced by a lateral nectary. In the box plot, the boxes represent the first and third quartile, the dark center lines represent the median values. The whiskers indicate minimal and maximal values. Nectar volumes were determined by measuring the diameter of each nectar droplet from wild-type (n=23) and *brcwinv4a-1* (n=43) flowers and using the formula for a sphere, V=4/3πr^3^. (C) Visualization of sucrose and glucose in nectar using a colorimetric assay. Nectar isolated from wild-type or *brcwinv4a-1* was diluted 500 or 50 times with water, respectively, prior to conducting the colorimetric assay. The amount of glucose content was determined with or without pretreatment of the diluted samples with commercial invertase. The quantified sucrose and glucose content is shown in Table 1.

**Table 1.** Nectar sugar content from lateral nectaries.

|  | Sugar (nmol/nectary) |  |
| --- | --- | --- |
|  | Sucrose | Glucose |
| *BrCWINV4A* / *BrCWINV4A* (Wild-type) | 1.3 ± 1.5 | 208.9 ± 38.6 |
| *brcwinv4a-1* / *BrCWINV4A* (hetero) | 0.9 ± 0.7 | 171.4 ± 19.1 |
| *brcwinv4a-1* / *brcwinv4a-1* (homo) | **17.3 ± 0.9\*\*\*** | **0.6 ± 0.17\*\*** |

A single replicate consisted of nectar collected from five independent flowers pooled into a single tube, with three replicates for collected for each genotype ( $N=3$ ). Bolded numbers indicate significant differences between wild-type and *brcwinv4a-1* (*homozygote*) according to according to a two-tailed Student's *t*-test (\*\* $P < 0.01$ ; \*\*\* $P < 0.001$ ). Standard error ( ± ) is indicated.

In addition to differences in nectar secretion patterns, the nectar droplets from *brcwinv4a-1* flowers appeared to be smaller than those of wild-type (Fig. 3A). Using image analysis and the formula for the volume of a sphere, we estimated the volume of nectar droplets secreted from the LNs of wild-type and *brcwinv4a-1* (Fig. 3B). The median volumes of lateral nectar droplets were 54.44 nL in wild-type and 7.7 nL in *brcwinv4a-1*. Considering that ∼29% of the mutant flowers produced no nectar, the total nectar output of *brcwinv4a-1* is estimated to be only ∼10% that of wild-type.

Since *atcwinv4a-1* flowers do not make nectar (Ruhlmann et al., 2010), it was surprising to find that a majority of *brcwinv4a-1* flowers do. Given that invertases hydrolyze sucrose into glucose and fructose, we examined the sugar content of *brcwinv4a-1* floral nectar. Fig. 3C shows the enzymatic visualization of glucose content in diluted nectar samples. Since this colorimetric assay only detects D-glucose, sucrose content was determined by adding exogenous, commercially available invertase to diluted nectar samples prior to the reaction step. Assays of wild-type nectar produced a bright pink color with or without pretreatment with exogenous invertase, indicating that the nectar has high glucose content. On the other hand, enzymatic reactions with *brcwinv4a-1* nectar without invertase added remained mostly colorless, but addition of invertase prior to the assay led to an intense pink color, indicating that there is high sucrose and little to no glucose in *brcwinv4a-1* nectar. This colorimetric assay was subsequently quantified with standard curves and confirmed that both wild-type and heterozygous (*BrCWINV4A-1/brcwinv4a-1*) flowers secrete a hexose-rich nectar with a comparable amount of total sugar (Table 1). Conversely, *brcwinv4a-1* secreted a sucrose-rich nectar, albeit with significantly less total sugar (Table 1).

We subsequently analyzed the sugar content in the LNs (tissue minus nectar) of both of wild-type and *brcwinv4a-1* (Fig. 4A). Predictably, *brcwinv4a-1* LNs had significantly higher sucrose (11.9 ± 0.4 nmol/nectar) and lower glucose content (2.7 ± 0.3 nmol/nectar) relative to wild-type (8.2 ± 0.7 nmol/nectar of sucrose and 9.0 ± 0.6 nmol/nectar of glucose) (Fig. 4A). In the case of median nectaries, sucrose content increased in the mutant, but glucose content was not significantly different (Supplemental Fig. S3). We also stained secretory nectary tissues (from open flowers) for the presence of starch using iodine-potassium iodide (IKI) (Fig. 4B). Wild-type and *brcwinv4a-1* similarly accumulated starch in the guard cells of LNs (Fig. 4B), but little starch was detectable inside the nectary parenchyma (Fig. 4B, bottom panels). Of note, the guard cells of nectary stomata are known to accumulate starch after the onset of nectar secretion (Ruhlmann et al., 2010).

**Figure 4.**
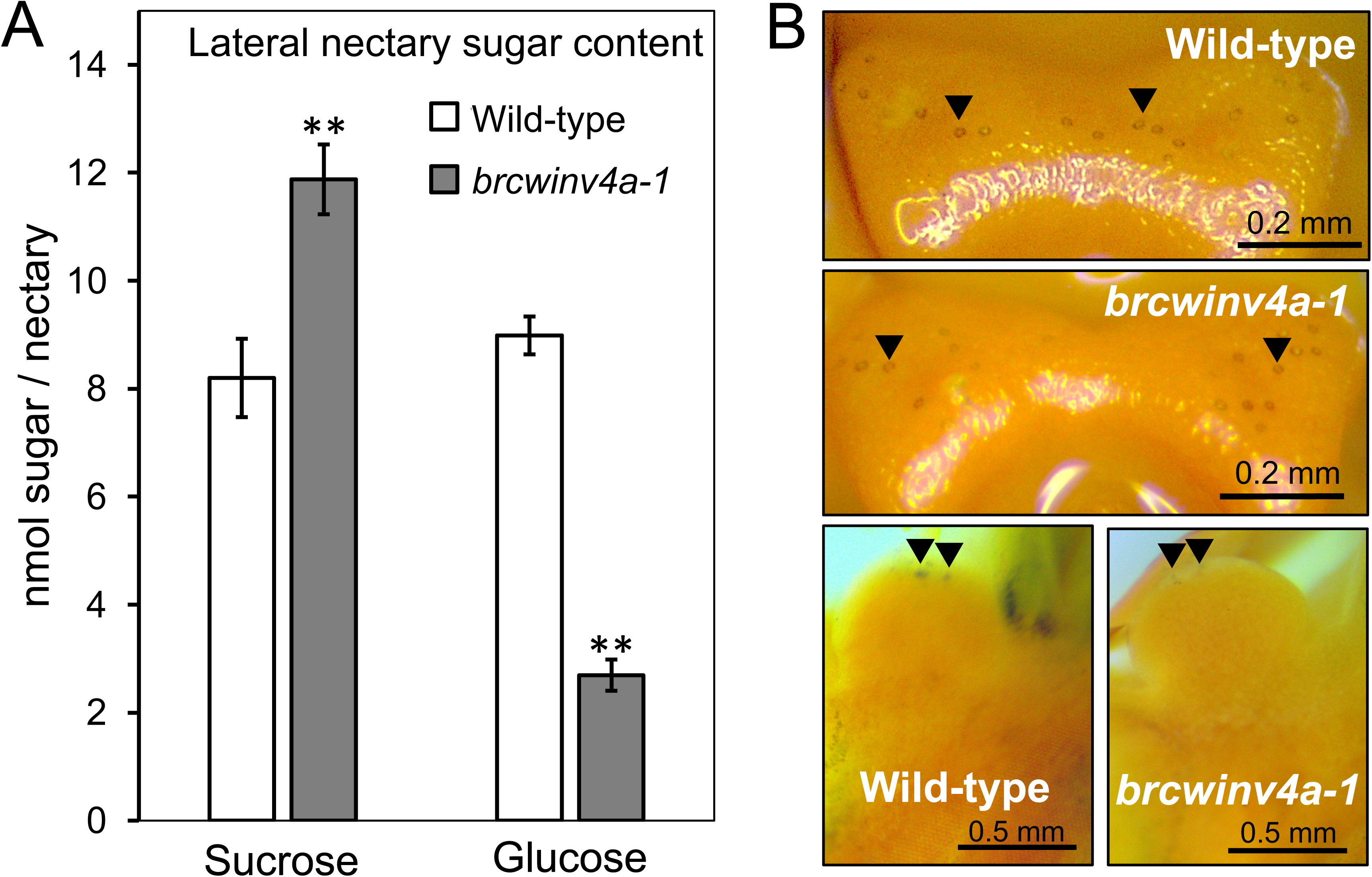
Sugar and starch content in the lateral nectaries of wild-type and *brcwinv4a-1*. (A) Sucrose and glucose content in lateral nectary tissues. Sucrose content was measured by pretreatment of samples with commercial invertase. Ten lateral nectaries collected from five independent flowers were pooled as a single sample for sugar assay. Bars represent averages of at least three independent biological and three technical replicates. Error bars show SE. Asterisks indicate significant differences between wild-type and *brcwinv4a-1* according to according to a Student’s *t*-test (**P < 0.001). (B) Starch accumulation in the lateral nectaries of wild-type and *brcwinv4a-1* as visualized with iodine-potassium iodide (IKI) stain. Upper two panels: intact nectaries with arrowheads pointing to representative stomata. Lower panels: starch staining of lateral nectaries after longitudinal sectioning. Arrowheads point to representative stomata.

Since the LNs of *brcwinv4a-1* produce less nectar than those of wild-type or heterozygous plants (Fig. 3), we investigated whether invertase activity in lateral nectaries can regulate nectar volume. For these experiments, ammonium heptamolybdate [(NH₄)_6_Mo_7_O_24_] was used because it is a potent inhibitor of plant invertases (Prado et al., 1979). To first demonstrate its inhibitory effect on the CWINV activity of *B. rapa* LNs, cell wall fractions extracted from wild-type LNs were incubated with ammonium heptamolybdate for 30 minutes at 30°C prior to measuring glucose released from exogenous sucrose (Figure 5A). Cell wall invertase activity was clearly inhibited by ammonium heptamolybdate at concentrations of 2 mM and higher, whereas the negative control treatment with ammonium chloride (NH₄Cl) did not inhibit CWINV activity. To then directly test the impact of this invertase inhibitor on nectar secretion, open flowers were first removed from wild-type inflorescences, with the cut stems (containing unopened flower buds) then being placed in 0.5x MS liquid medium (pH 5.2) containing either ammonium heptamolybdate or ammonium chloride (NH_4_Cl; as a control) for 1 to 3 days. After new flowers opened, we performed sampling sequentially and observed nectar production (Fig. 5B and Fig. 5C). Wild-type flowers treated with media containing NH_4_Cl had the same relative nectar producing capacity as mock-treated controls (Fig. 5B and 5C). Conversely, there was a dose-dependent reduction in nectar volume in ammonium heptamolybdate-treated flowers (Fig. 5).

**Figure 5.**
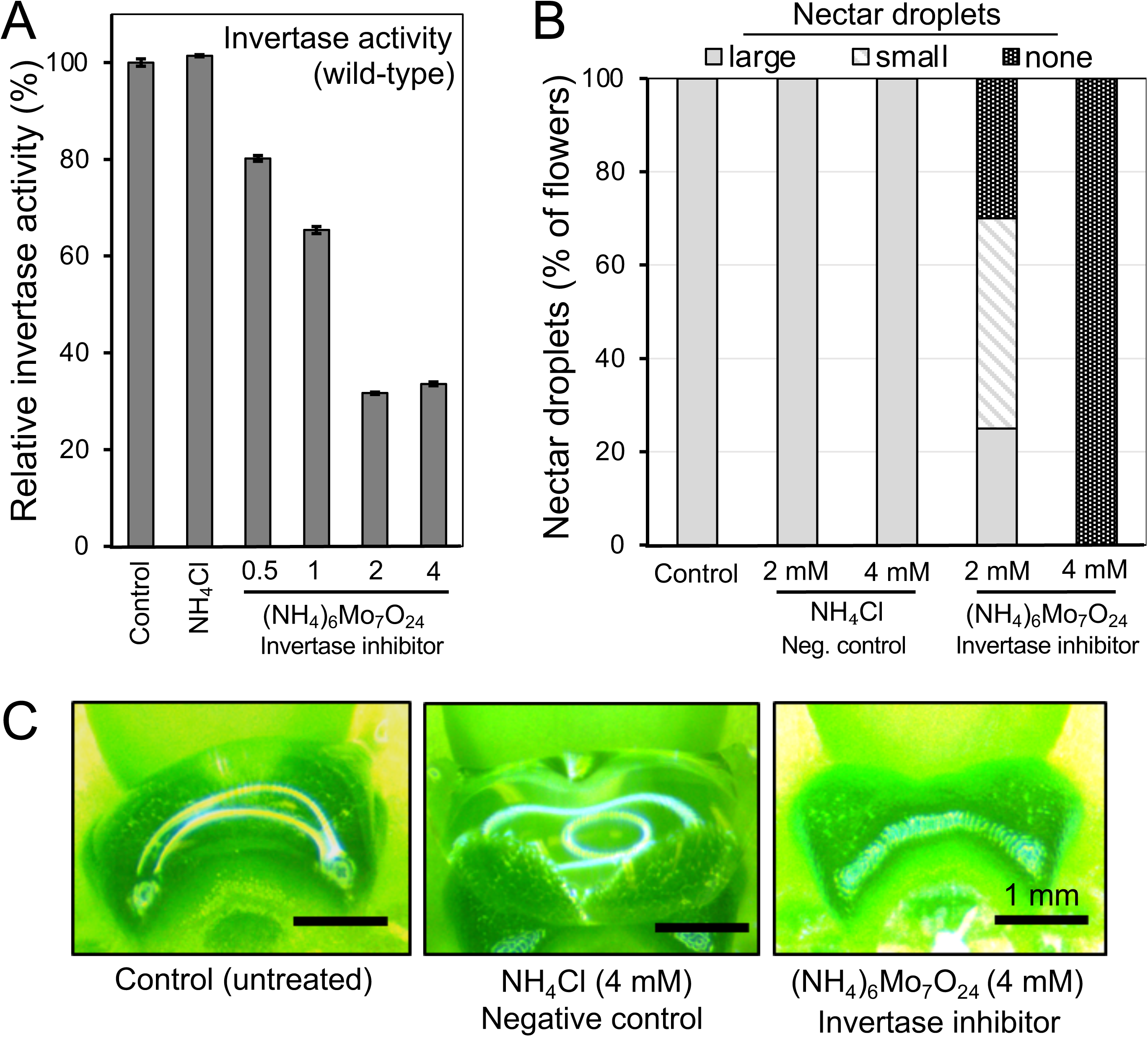
Effect of an invertase inhibitor, ammonium heptamolybdate, on nectar secretion of *Brassica rapa*. Wild-type inflorescence stems with open flowers removed (only flower buds remained) were incubated in liquid 0.5x MS media (pH 5.2) containing 1% (w/v) sucrose with or without ammonium chloride (NH_4_Cl) or varying levels of ammonium heptamolybdate [(NH_4_)_6_Mo_7_O_24_]. The inflorescence stems were incubated under high humidity conditions for 1-3 days and then observed for nectar secretion in newly opened flowers (post-treatment). (A) Relative cell wall invertase activity in controls, mock-treated (NH_4_Cl), and inhibitor-treated samples. (B) Percent nectar secretion phenotypes after inhibitor treatment with 0, 2, and 4 mM ammonium heptamolybate. Per nectary, these secretion phenotypes were classified as ‘large’ (one, large nectar droplet); ‘small’ (two small drops on the nectary surface), and ‘none’ (no nectar present), with the following replicates: mock control, n = 22; ammonium chloride [2 mM (n=16), 4 mM (n=16)]; ammonium heptamolybdate [2 mM (n=20), 4 mM (n=20)]. (C) Example images from treated flowers from A and B. Scale bars are 1 mm.

To test the field performance of *brcwinv4a-1*, both in terms of pollinator visitation and seed yield, we placed potted wild-type and *brcwinv4a-1* plants in the field at the initiation of flowering. Wild-type flowers were cumulatively visited by nearly two-fold more insects than those of *brcwinv4a-1* (Fig. 6), especially by honeybees and soldier beetles. However, *brcwinv4a-1* and wild-type had a similar number of visits by small carpenter bees, flies, and other bees and beetles. Wild-type plants also produced ∼30% more seed per plant than *brcwinv4a-1* (*P* = 0.00913, Table 2), but total seed weight per plant was not significantly different (*P* = 0.112). This lack of a significant difference in total seed weight produced per plant was due to the fact that *brcwinv4a-1* seeds were significantly larger (Area, *P* = 1.45 x 10^-14^) and weighed ∼17% more than those of wild-type (*P* = 0.00268, Table 2).

**Figure 6.**
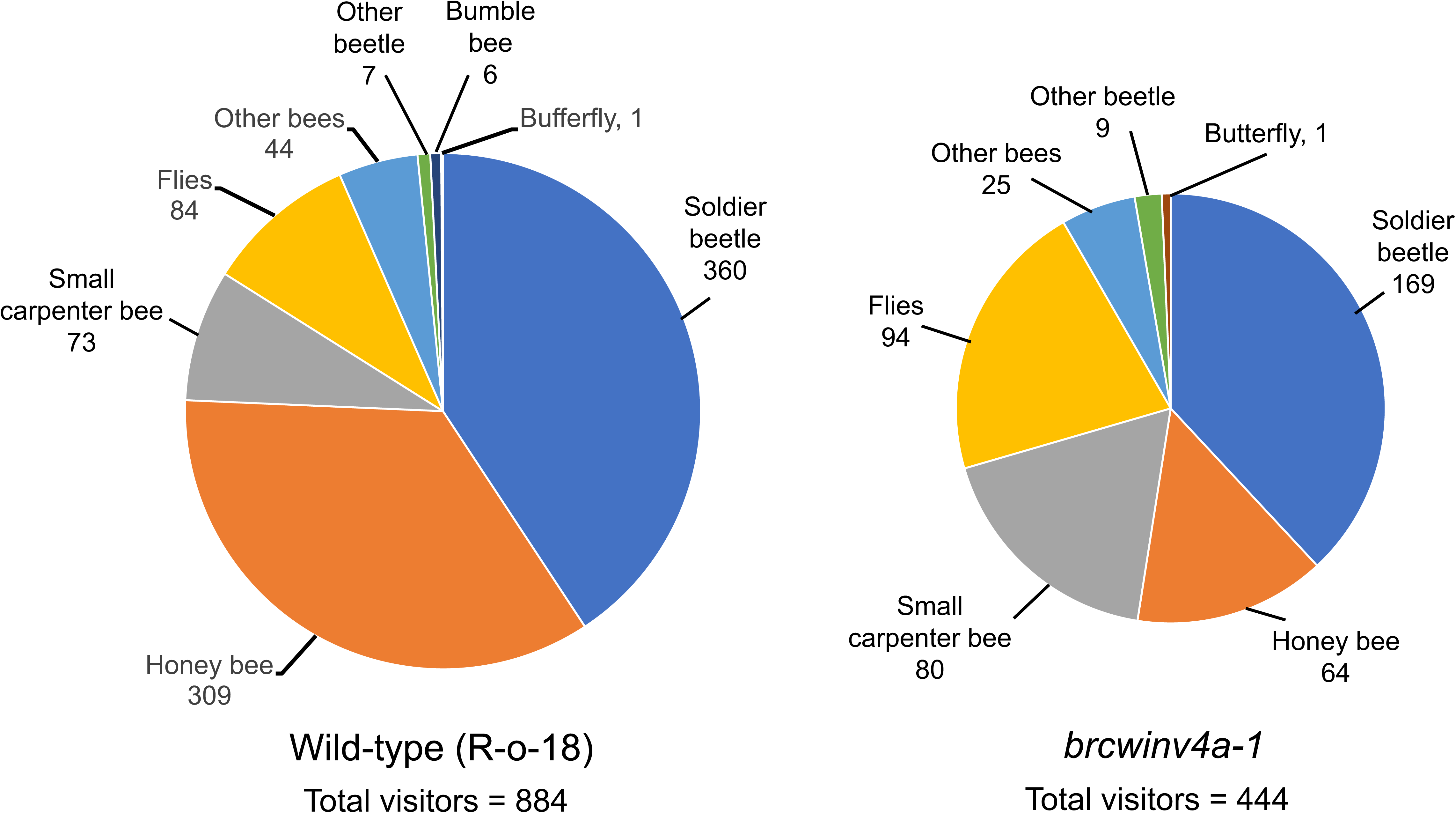
Pollinator visitation to wild-type and *brcwinv4a-1* flowers. The type and number of insect visitors to wild-type and *brcwinv4a-1* flowers are indicated as observed in the Minnesota Agricultural Experiment Station of the University of Minnesota (Falcon Heights, MN, USA). Twenty-two plants for each genotype were placed in rows of shallow trays separated by 5 m. Insect visitation was monitored for 5 minutes for each genotype (n = 42 times for each genotype) as described in the Materials and Methods.

**Table 2.** Seed size and yield in brwinv4a-1 and wild-type R-o-18.

|  |  | R-o-18<br>(n = 11) | *brcwinv4a-1*<br>(n = 12) | P-value |
| --- | --- | --- | --- | --- |
| **Seed yield** | Total seed weight per plant (g) | 6.96 ± 0.56 | 5.71 ± 1.65 | 0.112 |
|  | 1000 seed weight (mg) | 14.09 ± 0.59 | 16.59 ± 0.15 | **2.68 × 10<sup>-3</sup>** |
|  | Estimated total seed number per plant | 2,443 ± 195 | 1,731 ± 499 | **9.13 × 10<sup>-3</sup>** |
| **Seed size** | Width (mm) | 1.68 ± 0.08 | 1.82 ± 0.07 | **8.49 × 10<sup>-13</sup>** |
|  | Length (mm) | 1.86 ± 0.08 | 2.03 ± 0.07 | **1.22 × 10<sup>-15</sup>** |
|  | Area (mm<sup>2</sup>) | 2.17 ± 0.20 | 2.60 ± 0.18 | **1.45 × 10<sup>-14</sup>** |
|  | Circularity | 1.16 ± 0.01 | 1.15 ± 0.01 | **1.74 × 10<sup>-3</sup>** |

## DISCUSSION

In this study we aimed to refine our understanding of nectar synthesis through the study of cell wall invertases expressed in *B. rapa* in nectaries. Specially, we found that while several cell wall invertases are expressed in *B. rapa* nectaries (Fig. 1 & 2A), knockout of a single a single gene, *BrCWINV4A* (Fig. 2B), leads to greatly reduced cell wall invertase activity (Fig. 2C). These results indicate that BrCWINV4A is the primary apoplastic invertase in *B. rapa* nectaries; however, unlike Arabidopsis *atcwinv4* mutants, ∼71% of *brcwinv4a-1* flowers still secrete nectar (Fig. 3A). Assuming glucose and fructose are roughly equimolar in nectar (being the hydrolysis products of sucrose), *brcwinv4a-1* nectar contains ∼93% sucrose and ∼7% hexoses (Fig. 3C, Table 1). However, this lack of sucrose hydrolysis led to significantly less volume and total sugar in the nectar of *brcwinv4a-1* relative to wild-type (Fig. 3B, Table 1). A concomitant elevation of sucrose and reduction of glucose was observed in nectary tissues (Fig. 4). These genetic results were further supported by the exogenous application of an invertase inhibitor, ammonium heptamolybdate, which decreased both nectary cell wall invertase activity and nectar volume in a dose-dependent manner (Fig. 5). In summary, we provide genetic and biochemical evidence that modulation of CWINV activity can regulate both nectar sugar composition and volume in *B. rapa*.

These results cumulatively indicate that BrCWINV4A is not only essential for producing a hexose-rich nectar, but also supports the recently developed model of nectar secretion summarized in Fig. 7A. Nectary structure in the Brassicaceae consists of three main components: (1) epidermis with stomata, (2) parenchyma cells, and (3) phloem sieve tubes (Davis et al., 1986; Denisow et al., 2016) (Fig. 7). Sucrose is the main sugar in the phloem sap of most plants, including the Brassicaceae (Lohaus and Schwerdtfeger, 2014; Bertazzini and Forlani, 2016). Phloem sap is unloaded as a ‘pre-nectar’ component from the sieve tube elements in the nectary parenchyma and is thought to either (1) be immediately secreted into nectar, or (2) first stored as starch in immature flowers, followed by breakdown at anthesis and re-synthesized into sucrose (Pacini and Nepi, 2007; Vassilyev, 2010; Heil, 2011; Roy et al., 2017; Solhaug et al., 2019; Solhaug et al., 2019); however, there is some evidence for both processes occurring coincidentally (Solhaug et al., 2019). Histological and genetic evidence strongly suggests that Brassicaceae nectaries accumulate at least some starch in immature flowers and then convert it into sucrose shortly before the initiation of nectar secretion (Ren et al., 2007; Lin et al., 2014). For instance, knockdown of sucrose-phosphate synthases abolished nectar secretion in Arabidopsis (Lin et al., 2014). Regardless of the source of nectary sucrose, the most supported model of nectar secretion involves sucrose export into the nectary apoplast via the plasma membrane localized sucrose uniporter SWEET9 (Lin et al., 2014). Being a passive uniporter, SWEET9’s transport activity is fully dependent on the presence of a sucrose concentration gradient (Lin et al., 2014). Upon export into the apoplast, sucrose is hydrolyzed into glucose and fructose by the cell wall localized CWINV4, which maintains a high intracellular-to-extracellular sucrose gradient and also creates a negative water potential causing water to move to the apoplast. Finally, the sugars and water accumulated into the apoplast are secreted as a nectar drop through the opened stomata on guard cells on the epidermis.

**Figure 7.**
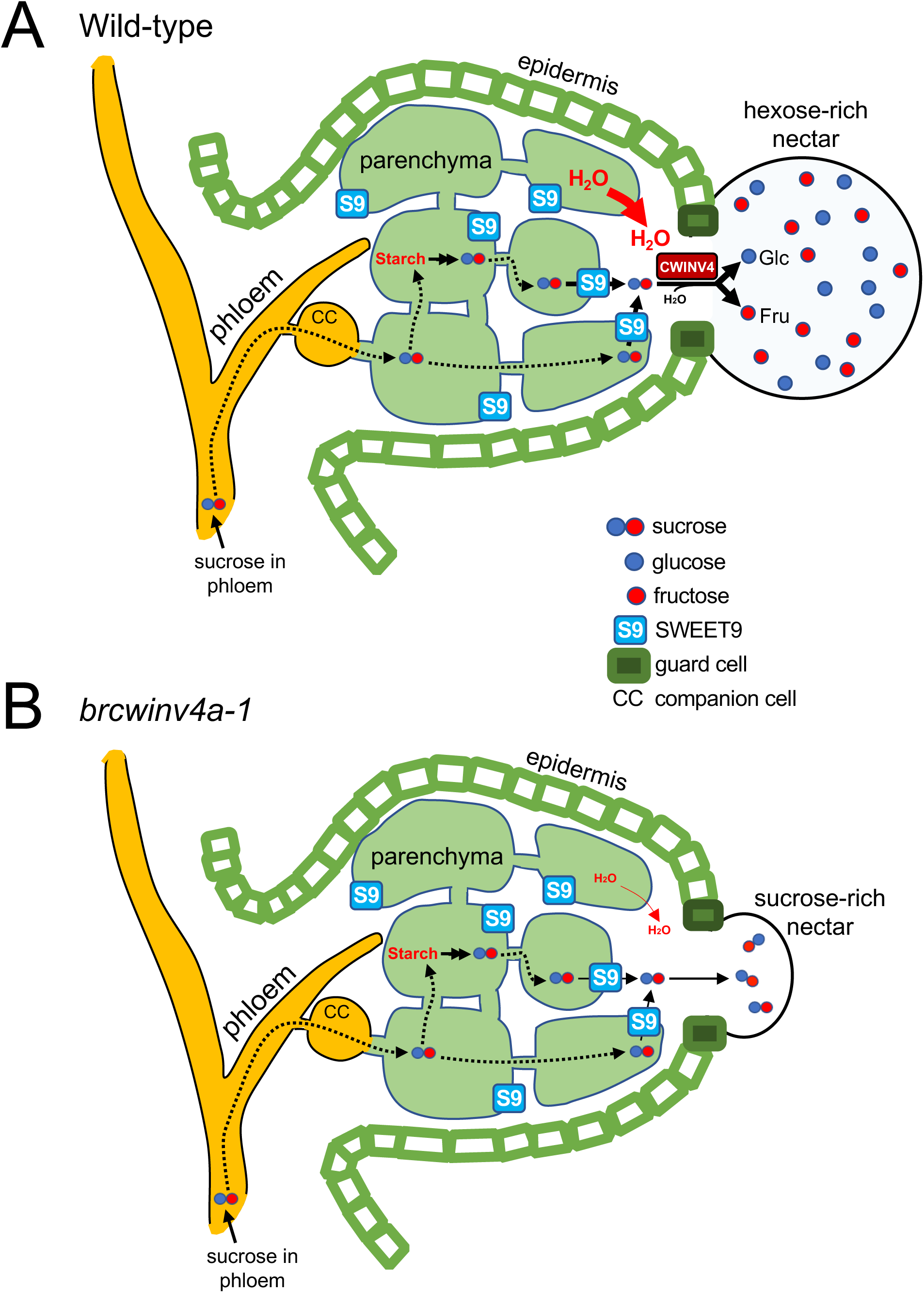
Simplified model of nectar secretion in a wild-type and *brcwinv4a-1* nectary. (A) In a wild-type flower, sucrose is delivered to this schematic nectary cross-section via the phloem and travels through the parenchyma where it is stored as starch. At anthesis starch is broken down and re-synthesized into sucrose by sucrose-phosphate synthase (SPS) and other enzymes. SWEET9 (S9), localized on the plasma membrane, then transports sucrose to the extracellular space in a purely concentration-dependent manner. Cell wall invertase activity (CWINV4) subsequently cleaves sucrose into hexoses to create a negative osmotic potential and a nectar droplet exiting via open stomates. Model based off of Kram and Carter, 2009 and Lin et al., 2014. <u>Note: for simplicity, only one CWINV4 is shown, although promoter::GUS studies suggest it is evenly distributed throughout the nectary parenchyma.</u> (B) In the absence of BrCWINV4 activity (i.e. in *brcwinv4a-1*), sucrose exported by SWEET9 is no longer hydrolyzed in the apoplast, thereby greatly reducing both the intracellular-to-extracellular sucrose concentration gradient and the osmotic potential for water flow out of nectary parenchyma cells, resulting in a smaller, sucrose-rich nectar droplet.

Within the context of this model of nectar secretion, loss of CWINV4 activity (Fig. 7B) would greatly decrease the intracellular-to-extracellular sucrose ratio in the nectary parenchyma, thereby diminishing the driving force for sucrose export via SWEET9. Additionally, without the subsequent hydrolysis of each sucrose into two hexoses, the osmotic potential required for nectar droplet formation would be strongly reduced. There is further genetic and biochemical evidence for cell wall invertases in regulating nectar sugar composition in both sunflower (*Helianthus annuus*) and squash (*Cucurbita pepo*) (Prasifka et al., 2018; Solhaug et al., 2019). For instance, most cultivated sunflower varieties have a hexose-rich nectar with little or no sucrose, but a recently described line with elevated sucrose (∼40% w/w) contained several polymorphisms in *HaCWINV2* (the closest homolog to *BrCWINV4* in the sunflower genome), which was associated with greatly reduced expression (Prasifka et al., 2018). However, Prasifka et al. (2018) did not report differences in nectar volume or cell wall invertase activity in sunflower nectaries. Conversely, squash naturally produces a sucrose-rich nectar with a large volume, which was explained by (1) very low expression and activity of cell wall invertases in its nectaries (Solhaug et al., 2019), and (2) the extremely high levels of intracellular sucrose (∼1M) being synthesized *de novo* in the nectary parenchyma (Solhaug et al., 2019). These two factors appear to facilitate the bulk flow of nectary sucrose out of the nectary and into the apoplast until a concentration equilibrium is reached on both sides of the plasma membrane of parenchyma cells containing SWEET9. It is likely that a similar phenomenon explains why *brcwinv4a* flowers secrete nectar when *atcwinv4* do not. Specifically, *B. rapa* nectaries are ∼10-fold larger than those of Arabidopsis, and thus likely have a higher capacity for intracellular sucrose generation to drive nectar droplet formation in the absence of CWINV activity. Alternatively, slight functional redundancy provided from BrCWINV2 and/or BrCWINV4B could explain the ability of *brcwinv4a* to secrete small volumes of a sucrose-rich nectar.

Nectar volume, concentration, and chemical composition all contribute to pollinator preference. For instance, some bees and butterflies often prefer nectars containing at least some sucrose (Wykes, 1952; Rusterholz and Erhardt, 1997; Rusterholz and Erhardt, 2000), and there tends to be positive correlation between nectar volume and pollinator visitation (Erickson, 1975; Cresswell, 1999). Thus, we predicted that the sucrose-rich nectar of *brcwinv4a-1* might be more palatable to some pollinators, while also acknowledging its low volume might make it less attractive for repeated visitation. In our limited study, wild-type flowers were visited much more heavily by honeybees and soldier beetles than those of *brcwinv4a-1* (Figure 6). While the ability to draw definitive conclusions will require further in-depth behavioral studies, these results strongly suggest that the potentially attractive sucrose-rich nectar of *brcwinv4a-1* cannot compensate for its relatively low volume, particularly as a reward for honeybees.

Perhaps unsurprisingly, wild-type plants produced significantly more seeds per plant than *brcwinv4a-1* (Table 2), likely due to increased pollinator visitation (Fig. 6). Even though *B. rapa* is not an obligate outcrosser, it has outcrossing rates of up to ∼40% in field conditions (Downey and Rimmer, 1993) and yields are greatly increased in the presence of bees and other pollinators (Mishra and Sharma, 1988; Bommarco et al., 2012). Although *brcwinv4a-1* produced less seed per plant, total seed weight was not significantly reduced relative to the wild-type, which was due to an increase in seed size and weight (Table 2). The larger seeds produced by *brcwinv4a-1* can possibly be explained by more energy being dedicated per seed (i.e., the same amount of total photosynthate as in wild-type spread over fewer seeds). An alternative, non-mutually exclusive explanation could be that *brcwinv4a-1* plants lose less carbon and energy through nectar production than wild-type. Indeed, up to 30% of total daily photosynthate can be dedicated to nectar production in some species (Pleasants and Chaplin, 1983), but it is currently unknown how much photosynthate goes towards nectar production in *B. rapa*. Although unlikely, we cannot exclude the possibility that secondary mutations might be responsible for the observed differences in insect visitation, seed set, and seed size between wild-type and *brcwinv4a-1*.

## CONCLUSIONS

Given the seemingly conserved mechanism of nectar secretion, at least within the eudicots (Lin et al., 2014; Solhaug et al., 2019; Chatt et al., 2021), our results strongly suggest that modulation of cell wall invertase activity can at least partially account for the observed differences in nectar sugar composition and volume, both within and between species. It will be particularly interesting to determine if a similar mechanism of nectar secretion holds true across more distant plant families and into monocots, as well as its role in shaping pollinator preference and plant fitness. This is especially true with the knowledge that nectaries have likely evolved multiple independent times [e.g. (Johnson et al., 2013)]. In a more applied context, the ability to control nectar sugar composition and volume in some crops could increase pollinator visitation and subsequent yield (Prasifka et al., 2018).

## MATERIAL AND METHODS

### Plant growth conditions

*Brassica rapa* (R-o-18) was the parental and wild-type line used in this study. Seeds were sterilized with a bleach solution (50% bleach, 0.05% Tween 20) for 8 min and washed three times with water prior to sowing. The sterilized seeds were grown with Berger BM2 germinating mix soil supplemented with Osmocote Flower and Vegetable Smart-release Plant Food. All plant growth was performed with a 16 hr light/8 hr dark cycle, photosynthetic photon flux of ∼200 µmol m^−2^ s^−1^ at 23-25°C. A TILLING (Targeting Induced Local Lesions in Genomes) mutant line (ji40447-b) was obtained from RevGenUK at the John Innes Centre (https://www.jic.ac.uk/research-impact/technology-platforms/genomic-services/reverse-genetics/) and the genotype was determined by sequencing of genomic PCR products with the following primer set: FW: CGGTTCAGCTTTCCGTGACCCG; RV: ACCGGGTGCTTACCTTTGACCCA. After four backcrosses (BC) to wild-type (R -O -18) and self-pollination, we used BC4F_1_, BC4F_2_ or BC4F_3_ plants for the experiments, unless otherwise noted. The sequences of additional primers used in this study can be found in Supplemental Table S1.

### Phylogenic analysis

The cell wall invertase amino acid sequences of *Brassica rapa* and Arabidopsis were obtained from the Brassica Database (BRAD, http://brassicadb.org) and TAIR (www.arabidopsis.org), respectively. The phylogenetic tree was conducted in MEGA X program (Kumar et al., 2018; Stecher et al., 2020) and generated using the Neighbor-Joining method (Saitou and Nei, 1987). The alignment was further optimized by manual inspection and curation. Seqboot was used for bootstrapping with 1,000 replicates. All ambiguous positions were removed for each sequence pair (pairwise deletion option). The evolutionary distances were computed using the JTT matrix-based method (Jones et al., 1992).

### RT-PCR Analyses

Plant tissues (leaf, stamen, calyx, petal, pistil, stamen, lateral nectary and median nectary) were frozen in liquid nitrogen after harvest and stored at -80°C. Total RNA was extracted using TRI Reagent (SIGMA) from the frozen *B. rapa* tissues. After DNase treatment with TURBO DNase-free^TM^ kit (Applied Biosystems), cDNA was synthesized from 1 µg total RNA with iScript^TM^ cDNA Synthesis Kit (Bio-Rad) These steps were carried out according to the manufacturer’s instructions. The cDNA template was used for PCR reaction with GoTaq Green Master Mix (Promega). The primers for RT-PCR are listed in Supplemental Table S1.

### Sucrose and glucose assay

Nectar drops in the lateral nectaries of a flower were collected by pipetting into 5 µl Milli-Q water. The samples of nectar collected from five independent flowers were gather into a tube and stored at -80 °C. For nectary samples, to remove nectar around nectary tissues, a flower without petal and calyx were washed into Milli-Q water twice, cut nectary tissues with a knife after wipe water around tissues and then stored at -80 °C. Sucrose and glucose contents in nectar and nectary tissues were determined with an Amplex Red Glucose/Glucose oxidase horseradish peroxidase method as previously (Bethke and Busse, 2008; Bender et al., 2012; Solhaug et al., 2019). Sucrose content was measured by pretreatment of samples with a commercial invertase (I9274, Sigma).

### Cell wall invertase activity assays

Cell wall invertase activity was assayed as previously described (Ruhlmann et al., 2010) with slight modifications. In brief, a single sample consisted of 20 lateral nectaries was collected from 10 flowers and stored at -80 °C before using the assay. The frozen tissue was grinded in an extraction buffer containing with 50 mM HEPES-NaOH (pH 8.0), 5 mM MgCl_2_, 2 mM EDTA, 1 mM MnCl_2_, 1 mM CaCl_2_, 1 mM DTT, and 0.1 mM PMSF and then centrifuged at 16,000×g for 10 min at 4 °C. The supernatant was discarded, and the extraction buffer added to the precipitation. After repeated 3 times, the precipitation was resuspended in 80 mM sodium acetate (pH 4.8). For the cell wall invertase assay, the extract was added to a sucrose solution (80 mM sodium acetate (pH 4.8), 5 mM sucrose) and then incubated at 30 °C for 30 min. The incubated sample were diluted by H_2_O and then produced glucose was measured using AmpRed Assay (Bethke and Busse, 2008; Bender et al., 2012; Solhaug et al., 2019).

### Measurement of nectar volume

We estimated one drop nectar size from two-drop type nectar on a lateral nectary from Stage 14–15 flower of wild type and *Brcwinv4A*. Images of two-drop type nectar were taken under a stereoscopic microscope (Amscope). The diameter of each nectar drop size from wild-type (n=23) and *brcwinv4a-1* (n=43) was measured using ImageJ software and calculated using the formula: V=4/3πr^3^. For large droplets of nectar on wild-type lateral nectary, we confirmed visual estimates of nectar droplet volume using a microcapillary tube (Drummond Microcaps**®** 1-000-001 Microliter).

### Invertase inhibitor treatment

The inflorescence stems of wild-type *Brasicca rapa* (R-o-18) plants we cut after removing all opened flowers, but leaving unopened buds intact. The cut inflorescence stems were quickly transferred into independently 2.0 mL tubes with 1.5 mL of autoclaved 0.5× Murashige and Skoog (MS) liquid media (pH5.2) containing 1% (w/v) sucrose. For inhibitor treatment, ammonium heptamolybdate [(NH_4_)_6_Mo_7_O_24_] and ammonium chloride (NH_4_Cl, control) were directly dissolved into liquid media of 0.5× MS media containing sucrose to reach 2 mM or 4 mM final concentration. A tube rack containing these tubes was placed in a tray lined with wet filter paper and covered with a transparent lid to maintain 100% humidity. The inflorescences were incubated in a plant growth room for 1 to 2 days. Opened flowers were subsequently collected and observed for nectar secretion under a stereoscopic microscope (Amscope), with a subset of nectaries being collected for cell wall invertase activity as described above.

### Starch staining and microscope observation

Open flowers (equivalent to Stage 14-15 in Arabidopsis) were fixed in glutaraldehyde solution (4% glutaraldehyde in 25 mM NaPO_4_, pH 7.3) as previously described (Thomas et al., 2017). After rinsing with ethanol, the flower samples were stained with Lugol’s iodine solution (Iodine Potassium Iodide 0.05M/0.1N, Fisher Science Education) as previously described (Ruhlmann et al., 2010).

### Pollinator visitation

Two independent sets of both wild-type and *brcwinv4a-1* seeds were planted in 8-inch diameter pots in the greenhouse (started on May 22, 2020 and June 6, 2020, respectively; n = 11 wild-type and *brcwinv4a-1* for each set (22 total plants of each genotype). Upon the initiation of flowering, the pots were moved to a field site (plot K-9) on the northeast corner of the Minnesota Agricultural Experiment Station of the University of Minnesota in Falcon Heights, Minnesota, USA (July 23, 2020 and August 5, 2020 for the two sets of plants described above, respectively). This plot was located <500 m from the University of Minnesota Bee Research Facility, which maintains an active apiary. The plants for each genotype were placed in rows of shallow trays separated by 5 m and watered by subirrigation daily. Insect visitation was monitored from July 28 – August 22, 2020 by counting floral visitors for 5 minutes for each genotype. All observations took place between 10 a. m - 3 p.m., but generally beginning at 12 p.m. The floral visitors for both wild-type and *brcwinv4-1* flowers were counted 42 times each and the data was binned and summed in aggregate.

### Seed analysis

For yield measurement and seed quality, we used the second set of the plants described for the pollinator visitation studies (planted on June 6, 2020 and moved to the field on August 5, 2020). Plants were allowed to finish flowering and have siliques mature in the field prior to returning them to the greenhouse to initiate drying (before siliques shattered). Seeds were manually harvested from siliques and dried at 37 °C for several days prior to measuring 200-seed weight. For the measurement of the seed size, we used a MARViN Seed Analyzer system (MARViTECH, Germany) as previously described (Esfahanian et al., 2021).

### Accession numbers

Arabidopsis invertases: AtCWINV1 = AT3G13790, AtCWINV2 = AT3G52600, AtCWINV3 = AT1G55120, AtCWINV4 = AT2G36190, AtCWINV5 = AT3G13784, AtCWINV6= AT5G11920, AtBFRUCT3 = AT1G62660, AtBFRUCT4 = AT1G12240. Brassica invertases: BrCWINV4A = BraA04g025740.3C, BrCWINV4B = BraA03g018900.3C, BrCWINV2 = BrA04g006780.3C. Additional *B. rapa* invertases used for phylogenetic analyses are noted by their accession numbers in Fig. 1.

## SUPPLEMENTAL MATERIAL

**Supplemental Table S1.** Oligonucleotide primers used in this study.

**Supplemental Figure S1.** Multiple protein sequencing alignment of *AtCWINV4*, *BraCWINV4A*, and *BraCWINV4B*.

**Supplemental Figure S2.** BrCWINV4B expression profiling and sequencing of PCR products.

**Supplemental Figure S3.** Multiple sequence alignment of a major *B. rapa* nectary cDNA sequence derived from ESTs (GQ146458) that encodes a cell wall invertase with the predicted translations of BraCWINV4A and BraCWINV4B.

**Supplemental Figure S4.** Examples of nectar secretion phenotypes in the F_2_ generation after the third backcross of plants genotyping as wild-type (*BrCWINV4A/BrCWINV4A*), heterozygous (*BrCWINV4A/brcwinv4a-1*) and homozygous (*brcwinv4a-1/brcwinv4a-1*). N > 10 for each genotype demonstrated consistent phenotypic patterns across genotypes.

## ACKNOWLEDGEMENTS

The authors thank Professors Donald Wyse and James Anderson, University of Minnesota Department of Agronomy and Plant Genetics, for arranging field space for the pollinator visitation studies and assisting with seed quality analysis, respectively. This work was funded by U.S. National Science Foundation grants IOS-0820730, IOS-1339246, IOS-2025297 to CJC.

## Abbreviations

CWINV: cell wall invertase
VINV: vacuolar invertase
SPS: sucrose-phosphate synthase
LN: lateral nectary
MN: median nectary

## LITERATURE CITED

1. Atmowidi T, Buchori D, Manuwoto S, Suryobroto B, Hidayat P (2007) Diversity of pollinator insects in relation to seed set of mustard (*Brassica rapa* L.: Cruciferae). HAYATI Journal of Biosciences 14: 155–161

2. Baker H, Baker I (1983) A brief historical review of chemistry of floral nectar. In BL Bentley, ed, The biology of nectaries. Columbia University Press, New York, pp 126–152

3. Bender R, Klinkenberg P, Jiang Z, Bauer B, Karypis G, Nguyen N, Perera MADN, Nikolau BJ, Carter CJ (2012) Functional genomics of nectar production in the Brassicaceae. Flora 207: 491–496

4. Berger S, Sinha AK, Roitsch T (2007) Plant physiology meets phytopathology: plant primary metabolism and plant-pathogen interactions. J Exp Bot 58: 4019–4026

5. Bertazzini M, Forlani G (2016) Intraspecific variability of floral nectar volume and composition in rapeseed (*Brassica napus* L. var. oleifera). Frontiers in Plant Science 7

6. Bethke PC, Busse JC (2008) Validation of a simple, colorimetric, microplate assay using Amplex Red for the determination of glucose and sucrose in potato tubers and other vegetables. Am J Potato Res 85: 414–421

7. Bommarco R, Marini L, Vaissiere BE (2012) Insect pollination enhances seed yield, quality, and market value in oilseed rape. Oecologia 169: 1025–1032

8. Chatt EC, Mahalim SN, Mohd-Fadzil NA, Roy R, Klinkenberg PM, Horner HT, Hampton M, Carter CJ, Nikolau BJ (2021) Nectar biosynthesis is conserved among floral and extrafloral nectaries. Plant Physiol doi.org/10.1093/plphys/kiab018

9. Cresswell JE (1999) The influence of nectar and pollen availability on pollen transfer by individual flowers of oil-seed rape (*Brassica napus*) when pollinated by bumblebees (*Bombus lapidarius*). Journal of Ecology 87: 670–677

10. Davis AR, Fowke LC, Sawhney VK, Low NH (1996) Floral nectar secretion and ploidy in *Brassica rapa* and *B. napus* (Brassicaceae) II. Quantified variability of nectary structure and function in rapid-cycling lines. Ann Bot 77: 223–234

11. Davis AR, Peterson RL, Shuel RW (1986) Anatomy and vasculature of the floral nectaries of *Brassica napus* (Brassicaceae). Can J Bot 64: 2508–2516

12. Davis AR, Pylatuik JD, Paradis JC, Low NH (1998) Nectar-carbohydrate production and composition vary in relation to nectary anatomy and location within individual flowers of several species of Brassicaceae. Planta 205: 305–318

13. Davis AR, Sawhney VK, Fowke LC, Low NL (1995) Floral secretion and ploidy in *Brassica rapa* and *B. napus* (Brassicaceae) .1. Nectary size and nectar carbohydrate production and composition. Apidologie 26: 534–534

14. Denisow B, Masierowska M, Anton S (2016) Floral nectar production and carbohydrate composition and the structure of receptacular nectaries in the invasive plant *Bunias orientalis* L. (Brassicaceae). Protoplasma 253: 1489–1501

15. Downey RK, Rimmer SR (1993) Agronomic improvement in oilseed *Brassicas*. Advances in Agronomy, Vol 50 50: 1–66

16. Erickson EH (1975) Variability of floral characteristics influences honey bee visitation to soybean blossoms. Crop Science 15: 767–771

17. Esfahanian M, Nazarenus TJ, Freund MM, McIntosh G, Phippen WB, Phippen ME, Durrett TP, Cahoon EB, Sedbrook JC (2021) Generating pennycress (*Thlaspi arvense*) seed triacylglycerols and acetyl-triacylglycerols containing medium-chain fatty acids. Frontiers in Energy Research 9: doi.org/10.3389/fenrg.2021.620118

18. Hampton M, Xu WW, Kram BW, Chambers EM, Ehrnriter JS, Gralewski JH, Joyal T, Carter CJ (2010) Identification of differential gene expression in *Brassica rapa* nectaries through expressed sequence tag analysis. PLoS ONE 5: doi.org/10.1371/Journal.Pone.0008782

19. Heil M (2011) Nectar: generation, regulation and ecological functions. Trends Plant Sci 16: 191–200

20. Johnson SD, Hobbhahn N, Bytebier B (2013) Ancestral deceit and labile evolution of nectar production in the African orchid genus Disa. Biology Letters 9: doi.org/10.1098/rsbl.2013.0500

21. Johnston JS, Pepper AE, Hall AE, Chen ZJ, Hodnett G, Drabek J, Lopez R, Price HJ (2005) Evolution of genome size in Brassicaceae. Annals of Botany 95: 229–235

22. Jones DT, Taylor WR, Thornton JM (1992) The rapid generation of mutation data matrices from protein sequences. Comput Appl Biosci 8: 275–282

23. Klein AM, Vaissiere BE, Cane JH, Steffan-Dewenter I, Cunningham SA, Kremen C, Tscharntke T (2007) Importance of pollinators in changing landscapes for world crops. Proceedings of the Royal Society B-Biological Sciences 274: 303–313

24. Kram BW, Carter CJ (2009) *Arabidopsis thaliana* as a model for functional nectary analysis. Sex Plant Reprod 22: 235–246

25. Kumar S, Stecher G, Li M, Knyaz C, Tamura K (2018) MEGA X: Molecular Evolutionary Genetics Analysis across computing platforms. Mol Biol Evol 35: 1547–1549

26. Lin IW, Sosso D, Chen LQ, Gase K, Kim SG, Kessler D, Klinkenberg PM, Gorder MK, Hou BH, Qu XQ, Carter CJ, Baldwin IT, Frommer WB (2014) Nectar secretion requires sucrose phosphate synthases and the sugar transporter SWEET9. Nature 508: 546–549

27. Lohaus G, Schwerdtfeger M (2014) Comparison of sugars, iridoid glycosides and amino acids in nectar and phloem sap of *Maurandya barclayana*, *Lophospermum erubescens*, and *Brassica napus*. Plos One 9:doi.org/10.1371/journal.pone.0087689

28. Masierowska ML (2003) Floral nectaries and nectar production in brown mustard (*Brassica juncea*) and white mustard (*Sinapis alba*) (Brassicaceae). Plant Systematics and Evolution 238: 97–107

29. Mishra R, Sharma S (1988) Growth regulators affect nectar-pollen production and insect foraging in Brassica seed crops. Curr Sci India 57: 1297–1299

30. Nicolson S, Thornburg R (2007) Nectar Chemistry. *In* SW Nicolson, M Nepi, E Pacini, eds, Nectaries and nectar. Springer, p 395

31. Ollerton J, Winfree R, Tarrant S (2011) How many flowering plants are pollinated by animals? Oikos 120: 321–326

32. Pacini E, Nepi M (2007) Nectar production and presentation. In SW Nicolson, M Nepi, E Pacini, eds, Nectaries and nectar. Springer, Dordrecht, pp 167–214

33. Paterson AH, Lan TH, Amasino R, Osborn TC, Quiros C (2001) Brassica genomics: a complement to, and early beneficiary of, the Arabidopsis sequence. Genome Biol 2: REVIEWS1011

34. Pierre J, Mesquida J, Marilleau R, Pham-Delegue MH, Renard M (1999) Nectar secretion in winter oilseed rape, Brassica napus - quantitative and qualitative variability among 71 genotypes. Plant Breeding 118: 471–476

35. Pleasants JM, Chaplin SJ (1983) Nectar production rates of *Asclepias quadrifolia*: causes and consequences of individual variation. Oecologia 59: 232–238

36. Prado FE, Sampietro AR, Vattuone MA (1979) Ammonium heptamolybdate, an inhibitor of plant invertases. Phytochemistry 18: 1799–1802

37. Prasifka JR, Mallinger RE, Portlas ZM, Hulke BS, Fugate KK, Paradis T, Hampton ME, Carter CJ (2018) Using nectar-related traits to enhance crop-pollinator interactions. Frontiers in Plant Science 9: doi.org/10.3389/Fpls.2018.00812

38. Ren G, Healy RA, Klyne AM, Horner HT, James MG, Thornburg RW (2007) Transient starch metabolism in ornamental tobacco floral nectaries regulates nectar composition and release. Plant Sci 173: 277–290

39. Roitsch T, Bittner M, Godt DE (1995) Induction of apoplastic invertase of *Chenopodium rubrum* by D-glucose and a glucose analog and tissue-specific expression suggest a role in sink-source regulation. Plant Physiology 108: 285–294

40. Roitsch T, Ehness R (2000) Regulation of source/sink relations by cytokinins. Plant Growth Regul 32: 359–367

41. Roy R, Schmitt AJ, Thomas JB, Carter CJ (2017) Review: Nectar biology: From molecules to ecosystems. Plant Sci 262: 148–164

42. Ruhlmann JM, Kram BW, Carter CJ (2010) CELL WALL INVERTASE 4 is required for nectar production in Arabidopsis. J Exp Bot 61: 395–404

43. Rusterholz HP, Erhardt A (1997) Preferences for nectar sugars in the peacock butterfly, *Inachis io*. Ecological Entomology 22: 220–224

44. Rusterholz HP, Erhardt A (2000) Can nectar properties explain sex-specific flower preferences in the Adonis blue butterfly *Lysandra bellargus*? Ecol. Entomol. 25: 81–90

45. Saitou N, Nei M (1987) The neighbor-joining method: a new method for reconstructing phylogenetic trees. Mol Biol Evol 4: 406–425

46. Sherson SM, Alford HL, Forbes SM, Wallace G, Smith SM (2003) Roles of cell-wall invertases and monosaccharide transporters in the growth and development of Arabidopsis. J Exp Bot 54: 525–531

47. Silva EM, Dean BB (2000) Effect of nectar composition and nectar concentration on honey bee (Hymenoptera: Apidae) visitations to hybrid onion flowers. J Econ Entomol 93: 1216–1221

48. Smyth DR, Bowman JL, Meyerowitz EM (1990) Early flower development in Arabidopsis. Plant Cell 2: 755–767

49. Solhaug E, Johnson E, Carter CJ (2019) Carbohydrate metabolism and signaling in squash nectaries and nectar throughout floral maturation. Plant Physiol 180: 1930–1946

50. Solhaug EM, Roy R, Chatt EC, Klinkenberg PM, Mohd-Fadzil N-A, Hampton M, Nikolau BJ, Carter CJ (2019) An integrated transcriptomics and metabolomics analysis of the *Cucurbita pepo* nectary implicates key modules of primary metabolism involved in nectar synthesis and secretion. Plant Direct 3: e00120

51. Stecher G, Tamura K, Kumar S (2020) Molecular Evolutionary Genetics Analysis (MEGA) for macOS. Mol Biol Evol 37: 1237–1239

52. Stephenson P, Baker D, Girin T, Perez A, Amoah S, King GJ, Ostergaard L (2010) A rich TILLING resource for studying gene function in *Brassica rapa*. BMC Plant Biol 10: 62

53. Sturm A (1999) Invertases. Primary structures, functions, and roles in plant development and sucrose partitioning. Plant Physiol 121: 1–7

54. Thomas JB, Hampton ME, Dorn KM, Marks MD, Carter CJ (2017) The pennycress (*Thlaspi arvense* L.) nectary: structural and transcriptomic characterization. Bmc Plant Biology 17

55. Town CD, Cheung F, Maiti R, Crabtree J, Haas BJ, Wortman JR, Hine EE, Althoff R, Arbogast TS, Tallon LJ, Vigouroux M, Trick M, Bancroft I (2006) Comparative genomics of *Brassica oleracea* and *Arabidopsis thaliana* reveal gene loss, fragmentation, and dispersal after polyploidy. Plant Cell 18: 1348–1359

56. Tymowska-Lalanne Z, Kreis M (1998) Expression of the *Arabidopsis thaliana* invertase gene family. Planta 207: 259–265

57. Tymowska-Lalanne Z, Kreis M (1998) The plant invertases: Physiology, biochemistry and molecular biology. Adv Bot Res 28: 71–117

58. Vassilyev AE (2010) On the mechanisms of nectar secretion: revisited. Annals of Botany 105: 349–354

59. Winter H, Huber SC (2000) Regulation of sucrose metabolism in higher plants: Localization and regulation of activity of key enzymes. Crit Rev Plant Sci 19: 31–67

60. Wykes GR (1952) The preferences of honeybees for solutions of various sugars which occur in nectar. Journal of Experimental Biology 29: 511–519

